# Evolutionary branching of host resistance induced by density-dependent mortality

**DOI:** 10.1101/410589

**Authors:** Jian Zu, Shuting Fu, Miaolei Li, Yuexi Gu

## Abstract

This study explores the evolutionary dynamics of host resistance in a susceptible-infected model with density-dependent mortality. We assume that the resistant ability of susceptible host will adaptively evolve, a different type of host differs in its susceptibility to infection, but the resistance to a pathogen involves a cost such that a less susceptible host results in a lower birth rate. By using the methods of adaptive dynamics and critical function analysis, we find that the evolutionary outcome relies mainly on the trade-off relationship between host resistance and its fertility. Firstly, we show that if the trade-off curve is globally concave, then a continuously stable strategy is predicted. In contrast, if the trade-off curve is weakly convex in the vicinity of singular strategy, then the evolutionary branching of host resistance is possible. Moreover, the bifurcation analysis shows that independent of the trade-off curve, the values of continuously stable strategy and evolutionary branching point will always increase as the demographic parameters increase. Secondly, after evolutionary branching in the host resistance has occurred, we examine the coevolutionary dynamics of the dimorphic host population and find that for a type of concave-convex-concave trade-off curve, the final evolutionary outcome may contain a relatively higher susceptible host and a relatively higher resistant host, which can continuously stably coexist on a long-term evolutionary timescale. Numerical simulation further shows that eventually the equilibrium population densities of the dimorphic susceptible host might be very close to each other. Finally, we find that for a type of sigmoidal trade-off curve, due to the high cost in terms of the birth rate, always the branch with higher resistance will go extinct, the eventual evolutionary outcome includes a monomorphic host with relatively lower resistance. Particularly, in this case we find that the evolution of costly host resistance may reduce the equilibrium population density of susceptible host, instead it may increase the equilibrium population density of infected host.

## Introduction

The host-pathogen interactions are ubiquitous in nature and hosts have evolved a diverse range of defence mechanisms in response to pathogens or parasites [1–15]. However, the underlying demographic and evolutionary factors are not well understood. An understanding of the evolutionary dynamics of host resistance is clearly necessary both to illustrate the host-pathogen interaction mechanisms and to manage effectively disease in a variety of application environments. Therefore, it is significant to investigate the evolutionary mechanism of host resistance in the light of ecological feedbacks [1–4, 7, 9, 10, 12, 13, 16].

In nature, in order to avoid to be infected by a pathogen, the resistant ability of host may adaptively evolve, but this is usually costly to acquire. In general, the higher resistant host may pay a cost in its other life-history traits, such as fertility. The existence of these costs is supported by both theoretical arguments and empirical evidences [1–4, 7, 9, 10, 12, 13, 17–27]. In order to understand the evolutionary mechanism of host resistance, in the past few decades, many different modeling approaches have been developed, such as locus-based method, quantitative genetic method and adaptive dynamics approach [4, 7, 9, 10, 12, 13, 19, 21, 28]. In particular, the method of adaptive dynamics allows us to examine the evolutionary factors that lead to different levels of host resistance. Based on the technique of adaptive dynamics [29–31], a number of different mathematical models have been proposed in order to understand how the host resistance evolves [1–4, 7, 9, 10, 12, 13, 19–21, 28, 32–38]. Antonovics and Thrall [2] and Bowers et al. [32] investigated a SI two-strain model. They assumed that the resistant host strain was less likely to become infected, but was more susceptible to crowding or had a lower birth rate. With the technique of analytical analysis and reciprocal invasion plot, they found that the evolution of host resistance depends mainly on the trade-off structure, highly susceptible strains can coexist with highly resistant strains. Boots and Haraguchi [19] extended this work to a multi-strain model. They assumed that different host strains differs in susceptibility to infection, with less susceptible host strains paying a cost leading to a lower intrinsic growth rate. Using the approach of adaptive dynamics and pairwise invasibility plot, they showed that the evolutionary outcome depends crucially on the shape of constraint function. With decelerating costly resistance, they found that evolutionary branching in the host resistance may occur and may lead to a dimorphic host with maximally resistant and no resistance. After branching in the host resistance occurs, based on an SIS (susceptible-infected-susceptible) model, Best et al. [13] showed that for sufficiently complex trade-off functions, the resulting invasion boundaries may form closed ‘oval’ areas of strategy coexistence. If there are no attracting singular strategies within this area, then the population may evolve out side of the coexistence region, leading to the extinction of one host strain. This phenomenon is also called as ‘evolutionary murder’, which has been observed in the studies of Geritz et al. [39]; Kisdi [40]; Parvinen [41]; Dercole and Rinaldi [42]; Best et al. [43]; Kisdi and Geritz [28].

In addition, the interaction between host and pathogen is commonly a coevolutionary process, both pathogen and host may adaptively evolve. Based on a SIS model, Restif and Koella [34] investigated the coevolutionary dynamics of host-pathogen interactions, they assumed that the transmission rate is determined by the traits of both host and pathogen and showed that the resistance reaches its maximum at an intermediate pathogen replication rate. Best et al. [9] also explored a coevolutionary model of a host-parasite system. They took the ecological feedbacks into consideration and found that evolutionary branching in the host resistance leads to parasites increasing their virulence, and small changes in the host resistance will drive large changes in the parasite virulence. Evolutionary branching in one species does not induce branching in the other.

However, in the previous studies, only the dynamics of susceptible host population includes density-dependence, the infected host population is not assumed to be subject to density-dependent mortality. More realistically, both susceptible host and infected host may be subject to density-dependent mortality. The natural death rate may be increased due to crowding through the density-dependence. Yet, there has been little discussion about how will the density-dependent mortality affect the evolution of host resistance. Moreover, in the previous studies, the trade-off function between host resistance and its intrinsic growth rate was assumed to be a specific form, and some of which were not a saturation function. But in reality, it is difficult to obtain the explicit formulation of trade-off function. In addition, after the branching has occurred in the host resistance, it is still unknown whether or not the coexistence of a dimorphic host can be maintained over evolutionary timescale and how does the finally evolutionary outcome depend on the shape and strength of trade-off relationship between the host resistance and its birth rate.

In this article, we assume that both susceptible host and infected host are subject to density-dependent mortality and a less susceptible host strain will pay a cost resulting in a lower birth rate. The trade-off relationship between host resistance and its fertility is assumed to be a monotonically increasing function and eventually converges to a saturated state. Based on an SI (susceptible-infected) model with density-dependent mortality, we will rigorously analyze the evolutionary invasion process of host resistance step by step. Both strictly mathematical proofs and numerical simulations are presented for each step of evolutionary invasion analysis. Specifically, the purpose of this study is aim to examine the following three questions. Firstly, under what demographic and evolutionary conditions a host population will change from monomorphism to dimorphism from the perspective of resistance. Using the method of critical function analysis [44–46, 48], we will identify the general properties of trade-off function that can induce the evolutionary branching in the host resistance and explore how will the shape and strength of trade-off relationship influence the evolutionary outcome. Besides, we will perform the bifurcation analysis of evolutionary dynamics and discuss the impact of demographic parameters on the evolutionary behaviour of host resistance. Secondly, under what demographic and evolutionary conditions the dimorphic susceptible host can continuously stably coexist on the long-term evolutionary timescale. Thirdly, whether the coevolution of dimorphism will result in the extinction of one strain. How complex a trade-off function must be in order for previously evolved dimorphism to be subject to evolutionary loss. Our main methods are based on the theory of adaptive dynamics and critical function analysis [12, 13, 26, 28–31, 44–52]. By using these approaches, we will clarify the interplay between ecological feedback and evolutionary feedback in the evolution of host resistance. Meanwhile, we will illustrate how the shape and strength of trade-off function are important in determining the level of resistance that evolves [10].

The rest of the paper is organized as follows. In the next section, we will present the population dynamics, evolutionary dynamics and critical function analysis. Afterwards, we summarize the demographic and evolutionary conditions that allow for a continuously stable strategy and evolutionary branching in the host resistance and discuss the influence of demographic parameters on the evolutionary outcome. In Section 4, we study the coevolutionary dynamics of a dimorphic susceptible host. Except for the strictly theoretical proof, numerical simulation results are also presented respectively in Sections 3 and 4 to illustrate the feasibility of our main results. A brief discussion is given at the end of this paper.

## Materials and Methods

We use the method of adaptive dynamics to analyze the evolutionary invasion process of host resistance step by step. Firstly, we will develop a population dynamics for an evolving susceptible host. Then based on this population model, the invasion fitness of mutant host and evolutionary model of host resistance will be derived. Finally, we use the method of critical function analysis to identify the evolutionarily singular strategy and its stability [44, 48, 50, 53, 54].

### Population dynamics

We consider a pathogen (a microparasite) that can cause lifelong infection of its host, and assume that the host population may develop resistance against this pathogen (i.e., the host population is able to defend against getting infected), but only at some cost [10]. Specifically, we assume that different host strains differ in how much they are resistant to the disease. A higher resistant host strain implies a lower rate of transmission the disease and thus a smaller value of transmission rate *β*(full resistance is achieved when *β* = 0). Therefore, the evolution of host resistance is transformed into the evolution of transmission rate *β*, but with an opposite relationship. Moreover, a higher resistant host strain is assumed to pay a cost in its other life-history trait, fertility, namely, a higher resistant host will result in a lower birth rate *b* such that *b*(*β*) is a continuous and monotonically increasing function with respect to *β* and eventually converges to a saturated state. In other words, there is a trade-off relationship between the transmission rate *β* and the birth rate of susceptible host *b*.

In addition, it is assumed that both the susceptible host population and infected host population are subject to density-dependent mortality *m*(*N)*, which is given by a linear function of total population density *N* = *S* + *I*, i.e.,

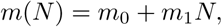

where *m*_0_ is the natural death rate of host, and *m*_1_ is a constant representing the strength of density-dependence. We also assume that infected individuals neither reproduce nor recover, which is a reasonable assumption for most invertebrate pathogens. In this way we focus on one possible route to disease resistance, that is, avoidance of infection by reduced susceptibility to the disease. Therefore, the population dynamics of a lifelong infection we have is given by

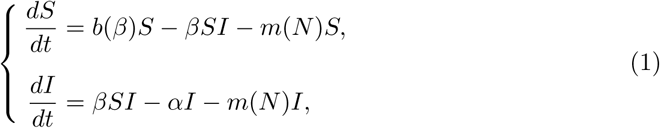

Where

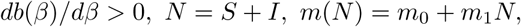

and *S* and *I* denotes respectively the population density of susceptible host and population density of infected host. *α* is the pathogen-induced mortality rate, also called the virulence. All of the parameters in model (1) are positive. The model is most applicable to invertebrate diseases and plant diseases [19].

Biologically, a convex trade-off curve *b*(*β*) means that decreasing *β* implies less and less decrease in *b*(*β*), that is to say, the cost increases slower than the benefit. In this case, we say there is a decelerating cost. In contrast, a concave trade-off curve *b*(*β*) means that decreasing *β* implies more and more increase in *b*(*β*), in other words, the cost increases faster than the benefit. In this case, we say there is an accelerating cost [10, 45, 53, 54].

For model (1), when

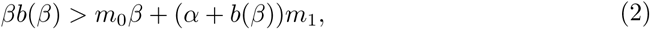

that is, the basic reproduction number

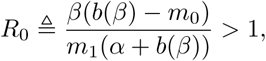

we obtain a strictly positive population equilibrium (*S*^*∗*^(*β*), *I*^*∗*^(*β*)), where

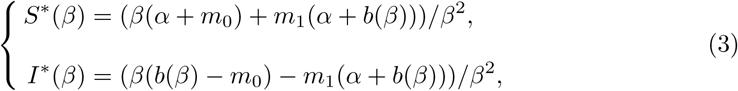

which is also globally asymptotically stable (see S1 Appendix for a detailed mathematical proof).

Next, based on model (1) and population equilibrium (*S*^*∗*^(*β*), *I*^*∗*^(*β*)), we will derive the evolutionary dynamics of host resistance.

### Invasion fitness of mutant host

We assume that only the host resistance (a type of phenotypic trait of susceptible host) can adaptively evolve, but the mutation in the host resistance is very small and the mutant host is rare. Before a mutant susceptible host appears, the resident hosts have reach to a population equilibrium state (*S*^*∗*^(*β*), *I*^*∗*^(*β*)). Therefore, the mutant host encounters resident population that is at the infected equilibrium state (*S*^*∗*^(*β*), *I*^*∗*^(*β*)) [30, 31, 49, 55].

When a mutant host with a slightly different resistance *β*_*m*_ enters into the resident population at a low density, the invasion fitness for mutant host [29] is given by

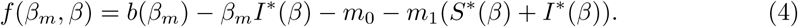

If *f* (*β*_*m*_, *β*) *>* 0, the population density of mutant host will increase, in other words, the mutant host can invade. By the results of Dercole and Rinaldi [42], Geritz et al. [56], Geritz [57] and Meszena et al. [58], we find that if the mutation in the host resistance is small, the mutant host is rare, and the transmission rate *β* is far from an evolutionarily singular strategy, then a successful invasion of mutant host will cause a trait substitution (see S2 Appendix for a detailed mathematical proof). Through successive invasions and replacements, the resistance of host will gradually evolve.

### Evolutionary dynamics of host resistance

The direction of such an evolutionary change is determined by the sign of selection gradient *g*(*β*), which is given by

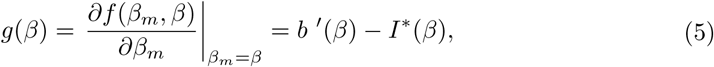

where *b* ^*′*^(*β*) denotes the derivative of *b*(*β*_*m*_) with respect to *β*_*m*_ evaluated at *β*_*m*_ = *β*.

By the results of Dieckmann and Law [30], if the mutation process is homogeneous and the mutation is small and rare, then the step by step evolution of host resistance can be approximated by the following equation

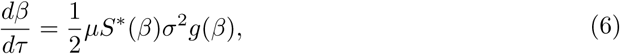

where time *τ* spans the evolutionary timescale and *μ* is the probability that a birth event in the susceptible host is a mutant, *S*^*∗*^(*β*) is the equilibrium population density of susceptible host, *σ*^2^ is the variance of phenotypic effect of mutation, and *g*(*β*) is the selection gradient described as in (5). The product 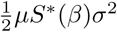 as a whole serves to scale the speed of evolutionary change [42, 51]. Equation (6) tells us how the expected value of host resistance *β* will change [53, 54]. Due to the uncertainty of the trade-off relationship *b*(*β*), we next use the method of critical function analysis to determine the ‘evolutionarily singular strategy’ and its stability.

### Critical function analysis

A strategy *β*^*∗*^ at which the selection gradient (5) equals to zero is called an ‘evolutionarily singular strategy’ [31, 51, 59–61]. From (5), we can see that a singular strategy *β*^*∗*^ should meet

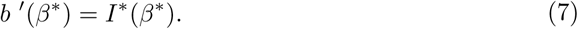

Because the trade-off function *b*(*β*) is uncertain, the singular strategy *β*^*∗*^ cannot be solved analytically. Therefore, we use the method of critical function analysis to distinguish the singular strategy and determine its stability [44, 48, 50, 53, 54].

Firstly, a critical function *b*_*crit*_(*β*) is a continuously differentiable function such that its slope is equal to the critical slope at all values of (*β, b*(*β*)) [44, 48, 50, 53, 54]. Thus, every strategy *β* on a critical function curve can be a singular strategy. From (3) and (7), we can see that the critical function is a solution of the following ordinary differential equation

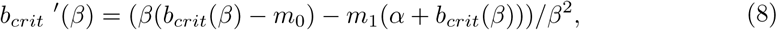

where a subscript is added to distinguish it from the trade-off function. Taking the derivative again with respect to *β* on both sides of the equation (8), we further get the curvature of critical function, which is given by

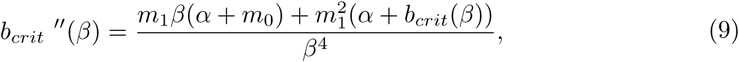

which is always positive, thus the curve of critical function is always convex. Generally speaking, given an arbitrary trade-off function, if the curve of trade-off function is tangential to a curve of critical function, then the point of tangency is an evolutionarily singular strategy [44, 48, 50, 53, 54].

Secondly, an evolutionarily singular strategy *β*^*∗*^ is convergence stable if a population with a nearby strategy can be invaded by a mutant host that is even closer to *β*^*∗*^ [31, 42, 51, 60, 62]. In other words, if *dg(β)/ dβ* |_*β = β*\*_ < 0, then the singular strategy *β*^*∗*^ is locally convergence stable. According to (5) and (9), we can see that if

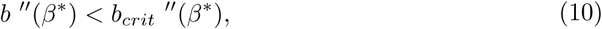

then the singular strategy *β*^*∗*^ is locally convergence stable [50]. Because the curve of critical function is always convex, the condition (10) implies that if the trade-off curve is less convex (or concave) than the curve of critical function in the vicinity of singular strategy *β*^*∗*^, then the singular strategy can be reachable [53, 54].

Thirdly, an evolutionarily singular strategy *β*^*∗*^ is locally evolutionarily stable if no nearby mutant can invade [31, 42, 51, 60, 62]. In other words, if *f* (*β*_*m*_, *β*^*∗*^) *<* 0 for all *β*_*m*_ *≠ β*^*∗*^ in a neighbourhood of *β*^*∗*^, then the singular strategy *β*^*∗*^ is locally evolutionarily stable. From (4), we can see that the singular strategy *β*^*∗*^ is locally evolutionarily stable if

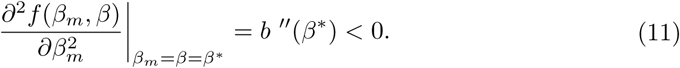

Therefore, if the trade-off curve is concave in the vicinity of *β*^*∗*^, then the singular strategy *β*^*∗*^ is locally evolutionarily stable. This implies that if there is an accelerating cost near the singular strategy *β*^*∗*^, then the singular strategy *β*^*∗*^ is evolutionarily stable [53, 54].

Given an arbitrary trade-off curve, the approach of critical function analysis gives a simple way to identify the evolutionarily singular strategy and its stability [44,48,50,53,54]. Combining the results of convergence stability and evolutionary stability, we obtain the following results.

## Results of monomorphic evolutionary dynamics

### Continuously stable strategy of host resistance

An evolutionarily singular strategy that is both convergence stable and evolutionarily stable is called a continuously stable strategy (CSS) [31, 44, 59]. A continuously stable strategy represents a finally evolutionary state of host resistance [31,59,61]. According to conditions (10) and (11), we obtain the following result.

**Theorem 1** *Assuming condition (2) holds*. *If one of the critical function b*_*crit*_(*β*) *is tangential to the trade-off curve b*(*β*) *at a strategy β*^*∗*^ *and b ″* (*β*^*∗*^) *<* 0, *then β*^*∗*^ *is an evolutionarily singular strategy of system (6) and it is continuously stable*.

From Theorem 1, we can see that if the trade-off curve is locally concave in the vicinity of *β*^*∗*^, then the singular strategy *β*^*∗*^ is a continuously stable strategy. This implies that if there is an accelerating cost near the singular strategy *β*^*∗*^, then natural selection will drive the evolution of host resistance toward *β*^*∗*^ and comes to a halt, namely, the singular strategy *β*^*∗*^ represents a final endpoint of the evolutionary process [31, 44, 59]. Thus the finally evolutionary outcome contains a single susceptible host with strategy *β*^*∗*^.

**Fig 1.**
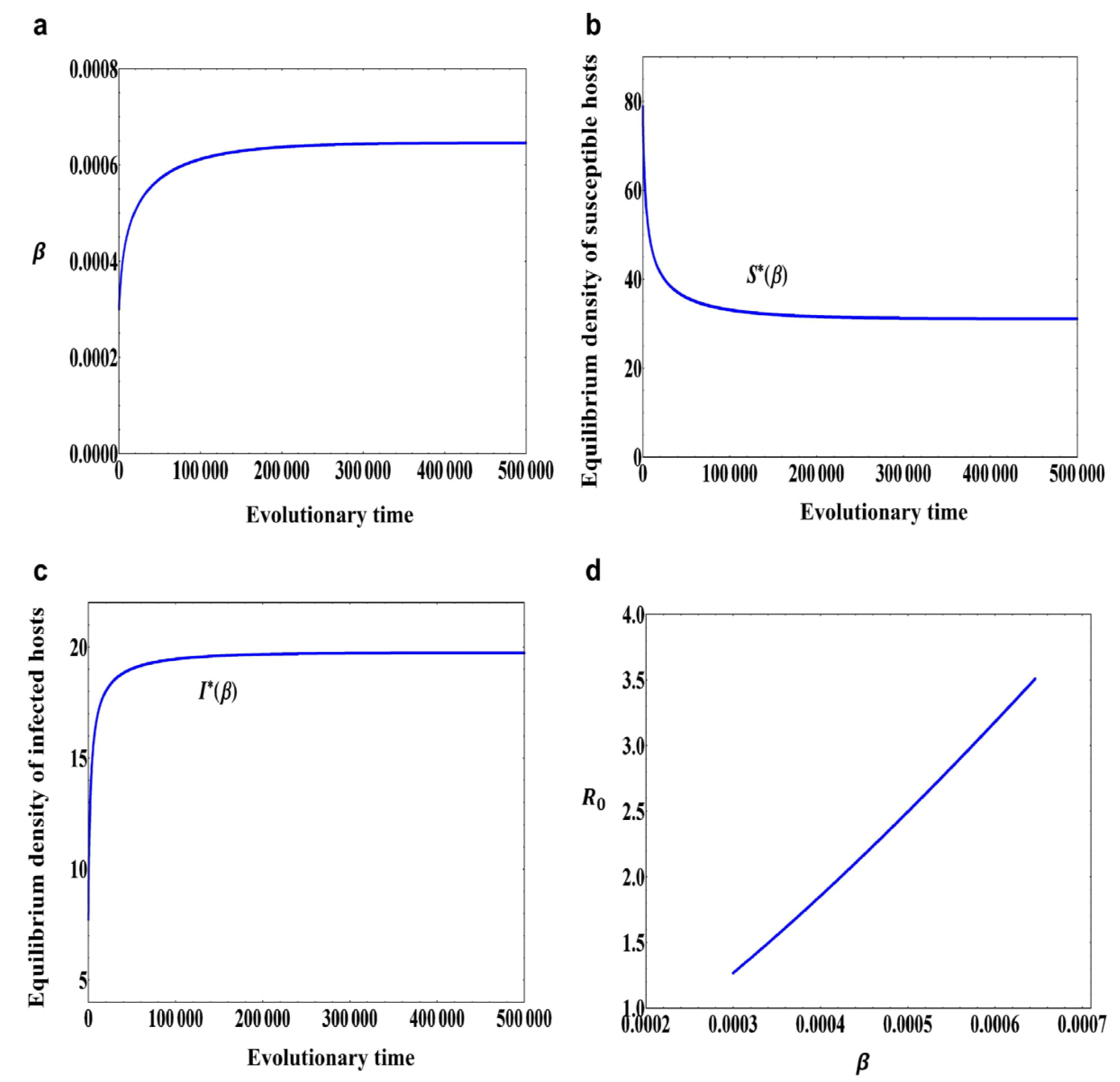
Continuously stable strategy of host resistance. (a) A trade-off curve *b*(*β*) described in (12) with *b*_0_ = 0.01, *b*_1_ = 2.0 and *b*_2_ = 0.1 and a tangential critical function curve *b*_*crit*_(*β*). The filled circle *β*^*∗*^ = 0.000646 is an evolutionarily singular strategy. The grey area indicates combinations of *β* and *b*(*β*) for which the positive population equilibrium, (*S*^*∗*^(*β*), *I*^*∗*^(*β*)), is globally asymptotically stable. (b) A pairwise invasibility plot (PIP) obtained with the trade-off function in (a). The grey areas indicate the mutant host can invade, while in green areas the mutant host becomes extinct. *β*^*∗*^ = 0.000646 is a continuously stable strategy. (c) An evolutionary time series plot of transmission rate *β* obtained through simulation of model (6) with initial condition *β*(0) = 0.0009. (d) Equilibrium population density of susceptible host *S*^*∗*^(*β*) when the transmission rate *β* evolves. (e) Equilibrium population density of infected host *I*^*∗*^(*β*) when the transmission rate *β* evolves. (f) The basic reproduction number *R*_0_ when the transmission rate *β* evolves. Other parameter values: *α* = 0.01, *m*_0_ = 0.005, *m*_1_ = 0.0001, *μ* = 0.01, *σ* = 0.00001.

As an example, the trade-off function is taken to be

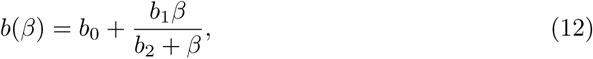

where *b*_0_ = 0.01, *b*_1_ = 2.0 and *b*_2_ = 0.1. In this case, *b*(*β*) is a monotonically increasing function and globally concave. Other parameter values are set to *α* = 0.01, *m*_0_ = 0.005, *m*_1_ = 0.0001, *μ* = 0.01, *σ* = 0.00001 (The values of these parameters and below are chosen such that they are basically consistent with the actual situation). In this case, we can see that the evolutionarily singular strategy *β*^*∗*^ in Fig.1a is both convergence stable and evolutionarily stable, that is, the evolutionarily singular strategy *β*^*∗*^ in Fig.1a is a continuously stable strategy (see also Fig.1b). From Fig.1c, we can see that if the initial resistance ability is relatively weak (*β*(0) = 0.0009), then the resistant ability of host will gradually increase. During the evolutionary process of host resistance, the equilibrium population density of susceptible host will also gradually increase and finally reaches to a saturation state (see Fig.1d), but the equilibrium population density of infected host will gradually decrease as the host resistance evolves (see Fig.1e). In this case, the evolution of host resistance results in a decrease of basic reproduction number *R*_0_ (see Fig.1f).

**Fig 2.**
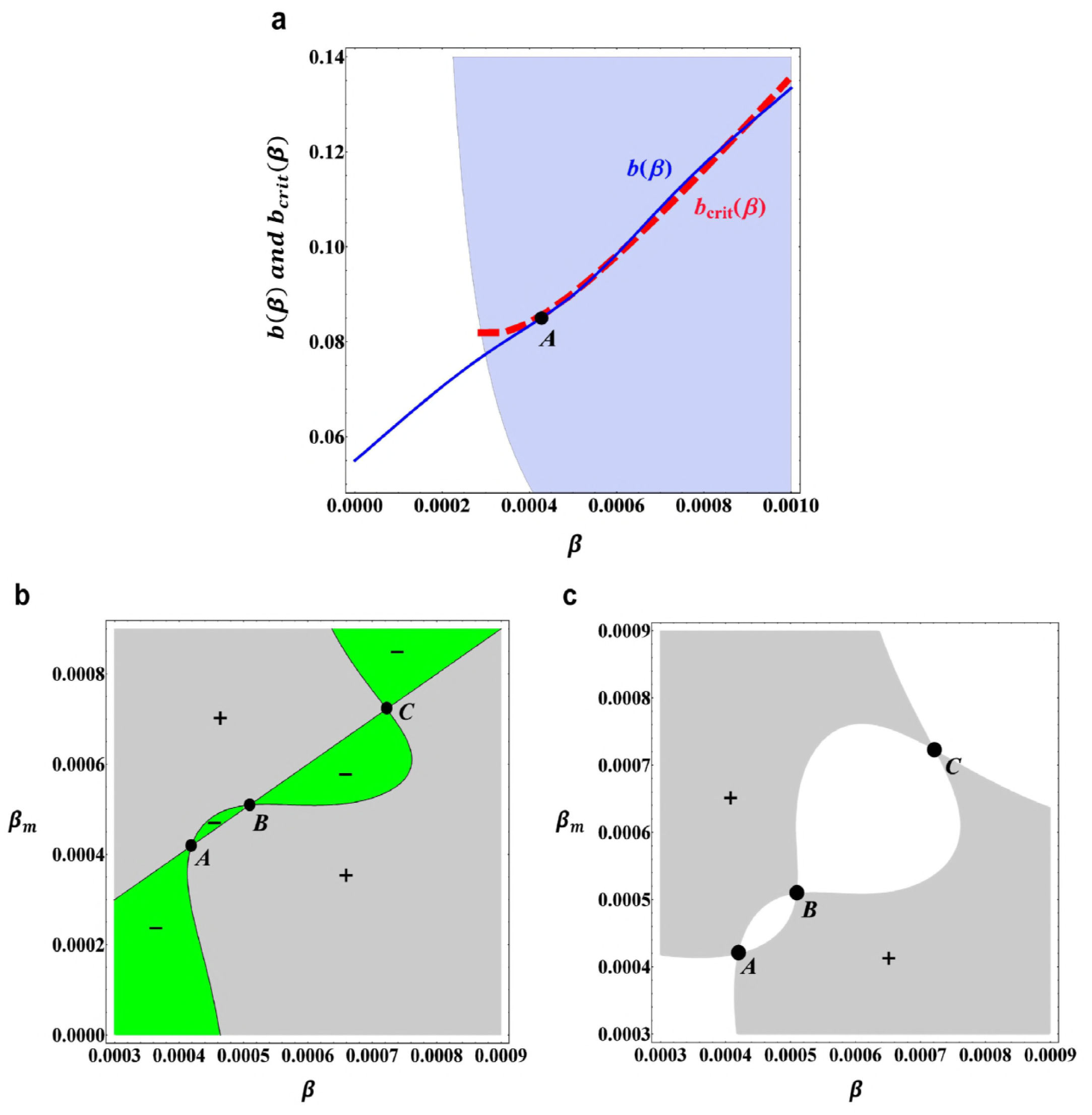
Continuously stable strategy of host resistance obtained with the same trade-off function as in Fig.1a, but with a different initial condition. (a) An evolutionary time series plot of transmission rate *β* obtained through simulation of model (6) with initial condition *β*(0) = 0.0003. (b) Equilibrium population density of susceptible host *S*^*∗*^(*β*) when the transmission rate *β* evolves. (c) Equilibrium population density of infected host *I*^*∗*^(*β*) when the transmission rate *β* evolves. (d) The basic reproduction number *R*_0_ when the transmission rate *β* evolves. Other parameter values are the same as in Fig.1.

However, if the initial resistant ability is relatively strong (*β*(0) = 0.0003), then due to the trade-off relationship between the birth rate and resistance, the evolution will reduce the resistant ability of susceptible host, the transmission rate *β* will gradually increase and converge to a saturation state (see Fig.2a). In this case, the equilibrium population density of susceptible host will gradually decrease (see Fig.2b), whereas the equilibrium population density of infected host will gradually increase and finally reaches to a saturation state (see Fig.2c). Particularly, in this case the evolution of host resistance leads to an increase of basic reproduction number *R*_0_ (see Fig.2d).

### Evolutionary branching of host resistance

If the evolutionarily singular strategy *β*^*∗*^ is convergence stable, but not evolutionarily stable, then by the results of Geritz et al. [31], Dercole and Rinaldi [42] and Geritz [57], the mutant host and resident host with strategies *β*_*m*_ and *β* close to *β*^*∗*^ can coexist and diverge in their resistant ability, that is to say, the susceptible host population will undergo evolutionary branching. Therefore, we obtain the following result.

**Theorem 2** *Assuming condition (2) holds*. *If one of the critical function b*_*crit*_(*β*) *is tangential to the trade-off curve b*(*β*) *at a strategy β*^*∗*^ *and* 0 *< b ″*(*β*^*∗*^) *< b*_*crit*_ *″*(*β*^*∗*^), *then β*^*∗*^ *is an evolutionarily singular strategy of system (6) and it is an evolutionary branching point (EBP)*.

From Theorem 2, we can see that near the singular strategy *β*^*∗*^ if the trade-off curve is less convex than the critical function curve, then this singular strategy *β*^*∗*^ is an evolutionary branching point (EBP). This means that in the vicinity of the singular strategy *β*^*∗*^ if there is a weakly decelerating cost, then evolutionary branching in the host resistance will occur.

**Fig 3.**
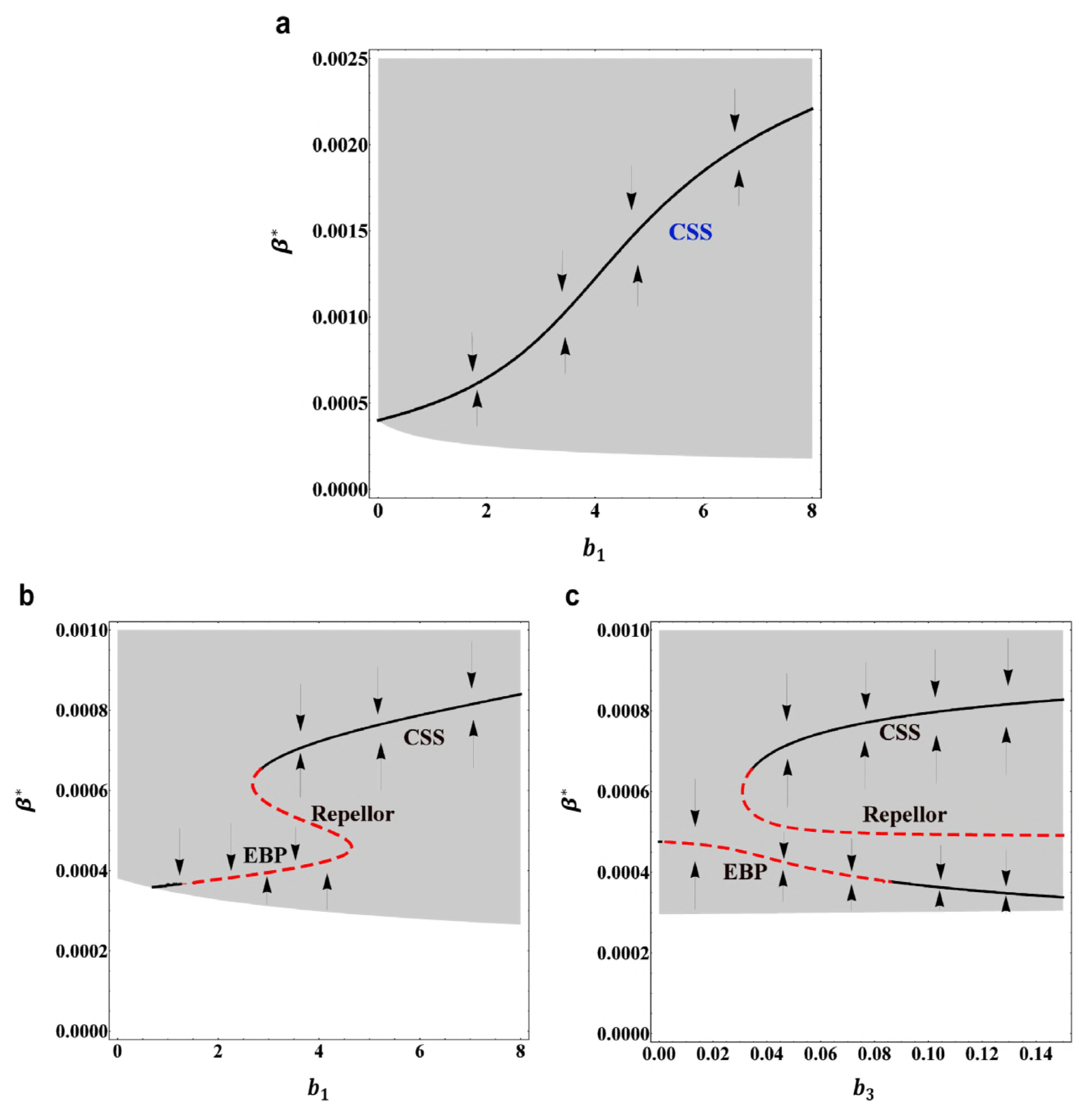
Evolutionary branching strategy of host resistance. (a) A trade-off curve *b*(*β*) described in (13) with *b*_0_ = 0.055, *b*_1_ = 4.0, *b*_2_ = 0.05, *b*_3_ = 0.05, *β*_0_ = 0.0005, *σ*_*c*_ = 0.0002 and a tangential critical function curve *b*_*crit*_(*β*). The filled circle *A* = 0.000419 is an evolutionarily singular strategy. The grey area indicates combinations of *β* and *b*(*β*) for which the positive population equilibrium, (*S*^*∗*^(*β*), *I*^*∗*^(*β*)), is globally asymptotically stable. (b) A pairwise invasibility plot (PIP) obtained with the trade-off function in (a). The grey regions indicate the mutant host can invade, while in green regions the mutant host becomes extinct. *A, B, C* are three evolutionarily singular strategies. (c) A mutual invasibility plot obtained with the trade-off function in (a). In the grey regions, both *f* (*β*_*m*_, *β*) *>* 0 and *f* (*β, β*_*m*_) *>* 0. Other parameter values: *α* = 0.075, *m*_0_ = 0.006, *m*_1_ = 0.00014, *μ* = 0.01, *σ* = 0.00001.

As an example, we take the following trade-off function [64]

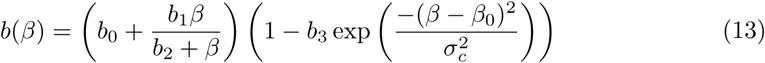

where *b*_0_ = 0.055, *b*_1_ = 4.0, *b*_2_ = 0.05, *b*_3_ = 0.05, *β*_0_ = 0.0005 and *σ*_*c*_ = 0.0002. In this case, *b*(*β*) is a monotonically increasing function and finally converges to a saturated state. Particularly, it is a concave-convex-concave trade-off curve. Other parameter values are set to *α* = 0.075, *m*_0_ = 0.006, *m*_1_ = 0.00014, *μ* = 0.01, *σ* = 0.00001. From Fig.3a, it can be seen that near the evolutionarily singular strategy *A* = 0.000419 the trade-off curve is less convex than the critical function curve, which implies that the evolutionarily singular strategy *A* = 0.000419 in Fig.3a is convergence stable but not evolutionarily stable. That is to say, the singular strategy *A* in Fig.3a is an evolutionary branching point (see also Fig.3b), close to this singular strategy *A* = 0.000419 the mutant host and resident host can coexist and diverge in their resistant ability (see Fig.3c), the evolutionary branching in the host resistance will occur. From Fig.3b, we can also see that if the initial resistant ability is relatively strong (i.e., with a smaller value of transmission rate), then the susceptible host will firstly evolve towards the smaller singular strategy *A*, and then branch into two different types. However, if the initial resistant ability is relatively weak (i.e., with a larger value of transmission rate), then the susceptible host will evolve towards a continuously stable strategy *C* and come to a halt.

### Results of bifurcation analysis

How likely does the continuously stable strategy of host resistance evolve or how likely does the evolutionary branching in the host resistance strategy occur? This can be observed through the bifurcation analysis of evolutionary dynamics. In this section, by using the method of numerical simulation, we examine how the evolutionary singular strategy and its stability will change as the shape and strength of the trade-off curve is changed.

**Fig 4.**
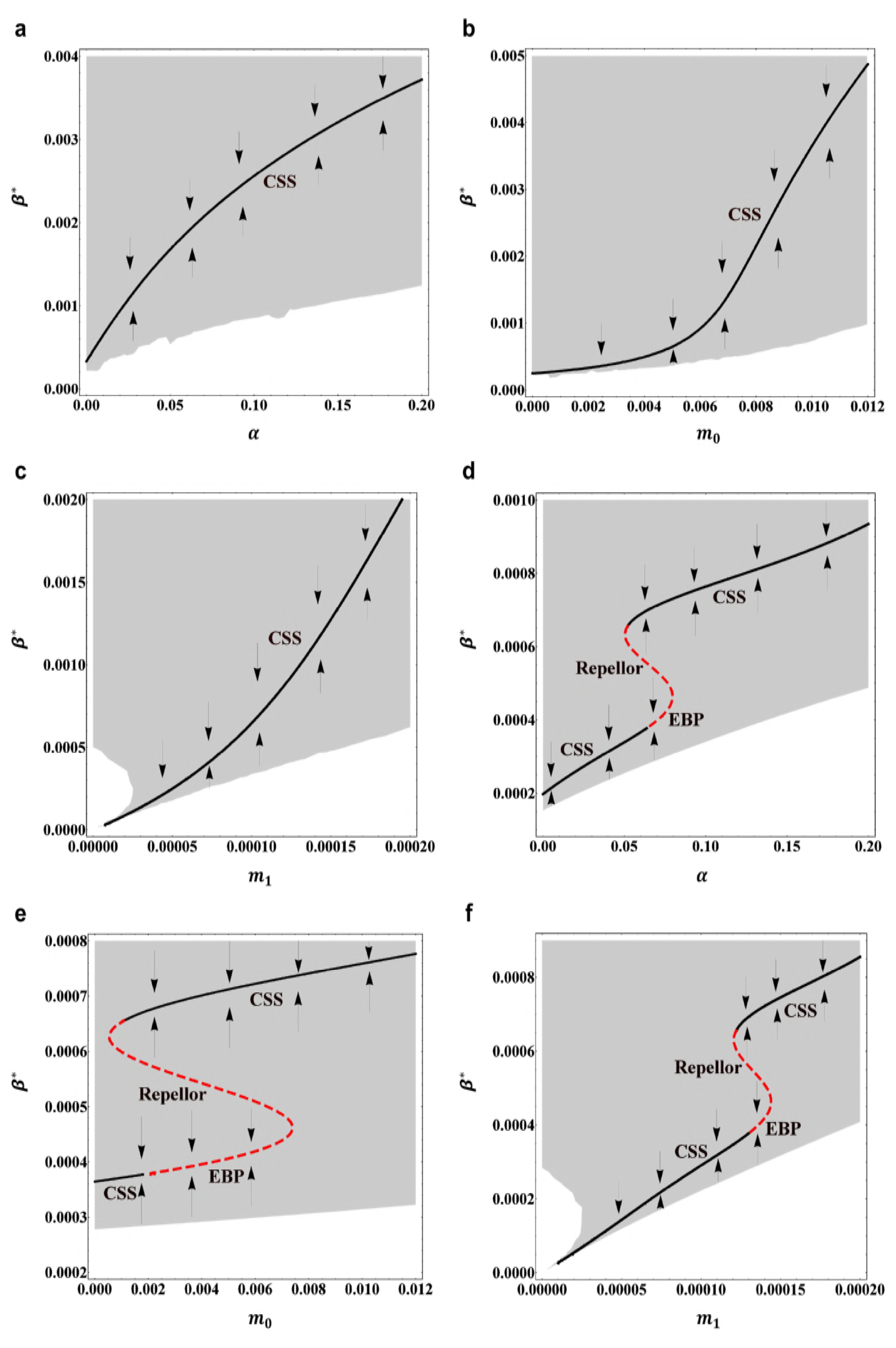
Bifurcation diagram of evolutionary dynamics when the parameter of tarde-off function is changed. (a) Evolutionarily singular strategy *β*^*∗*^ versus the maximum birth rate *b*_1_ when the trade-off function is the same as in Fig.1a. Other parameter values are the same as in Fig.1. Arrows indicate the direction of evolutionary change. *CSS* denotes the continuously stable strategy. (b) Evolutionarily singular strategy *β*^*∗*^ versus the maximum birth rate *b*_1_ when the trade-off function is the same as in Fig.3a. *CSS* denotes the continuously stable strategy, *EBP* denotes the evolutionary branching point. Other parameter values are the same as in Fig.3. (c) Evolutionarily singular strategy *β*^*∗*^ versus the trade-off strength *b*_3_ when the trade-off function is the same as in Fig.3a. Other parameter values are the same as in Fig.3. In (a), (b) and (c), the black solid curves indicate the evolutionarily singular strategies which are both convergence stable and evolutionarily stable; the red dashed curves indicate the evolutionarily singular strategies which are not evolutionarily stable. The grey region indicates combination of *β* and *b*(*β*) for which the positive population equilibrium is globally asymptotically stable.

For the globally concave trade-off function (12), the bifurcation diagram of evolutionarily singular strategy *β*^*∗*^ versus the maximum birth rate *b*_1_ is depicted in Fig.4a. Other parameter values are set to the same values as in Fig.1. We can see that if the trade-off function is globally concave, then the singular strategy *β*^*∗*^ is always convergence stable as well as evolutionarily stable, the natural selection will drive the evolution of host resistance towards *β*^*∗*^ and come to a halt. When the maximum birth rate *b*_1_ increases continuously from zero, the value of continuously stable strategy *β*^*∗*^ will gradually increase and finally reaches to a saturation state. This implies that when there is an accelerating cost, then decreasing the host resistance (i.e., increasing continuously stable strategy *β*^*∗*^, accordingly increasing the fertility of host) will benefit the susceptible host population. In particular, for the globally concave trade-off function, we can see that the evolutionary branching in the host resistance strategy cannot occur.

In the case of a concave-convex-concave trade-off function (13), the bifurcation diagram of evolutionarily singular strategy *β*^*∗*^ versus the maximum birth rate *b*_1_ is depicted in Fig.4b. Other parameter values are set to the same values as in Fig.3. In this case, if 0.69 *< b*_1_ *<* 8.0, then the trade-off function *b*(*β*) is monotonically increasing and finally converges to a saturation state. It can be seen that when 1.29 *< b*_1_ *<* 4.64, then the smaller singular strategy *β*^*∗*^ is convergence stable but not evolutionarily stable, close to this singular strategy, evolutionary branching in the host resistance strategy will occur. Particularly, we can see that when the maximum birth rate *b*_1_ gradually increases, the value of evolutionary branching point will also gradually increase. However, when 2.69 *< b*_1_ *<* 4.64 whether or not the evolutionary branching occurs also depends on the initial host resistance strategy, because in this case there also exists a continuously stable strategy for the host resistance.

Moreover, based on the trade-off function (13), Fig.4c depicts the bifurcation diagram of evolutionarily singular strategy *β*^*∗*^ versus the trade-off strength *b*_3_. Other parameter values are set to the same values as in Fig.3. In this case, when 0.0 *< b*_3_ *<* 0.15, the trade-off function *b*(*β*) is monotonically increasing and finally converges to a saturation state. From Fig.4c, we can see that when 0.0014 *< b*_3_ *<* 0.086, that is, for a moderately strong trade-off function, the smaller singular strategy *β*^*∗*^ is convergence stable but not evolutionarily stable, close to this singular strategy, evolutionary branching in the host resistance will occur. However, when 0.031 *< b*_3_ *<* 0.086, that is, the convexity of trade-off function becomes relatively strong, then whether evolutionary branching occurs or not also depends on the initial host resistance strategy, because in this case there also exists another continuously stable strategy (CSS) for the host resistance (see Fig.4c).

From Fig.4, we can see that given the environmental conditions and the functional form of trade-off relationship, there may exist a wide range of parameter values (such as *b*_1_ and *b*_3_) which can lead to evolutionary branching in the host resistance. Next, we further discuss the impact of changes of demographic parameters on the evolutionary behaviour of host resistance.

### Effect of demographic parameters

In this section, by using the method of implicit differentiation, we discuss how likely the evolutionary branching in the host resistance will occur independent of the trade-off function, and how changes in the demographic parameters will affect the evolutionary behaviour of host resistance.

First of all, we consider the impact of the strength of density-dependent mortality *m*_1_ on the continuously stable strategy (CSS) and evolutionary branching point (EBP) of host resistance. This can be determined by the sign of the derivative of *β*^*∗*^ with respect to *m*_1_. On the basis of the selection gradient *g*(*β, m*_1_) in (5), by implicitly differentiating the equation *g*(*β*^*∗*^, *m*_1_) = 0 with respect to *m*_1_, and treating *β*^*∗*^ as a function of *m*_1_ [53, 54], this gives

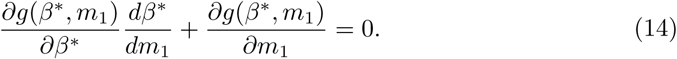

Using the definition of *g*(*β, m*_1_), we rewrite (14) as

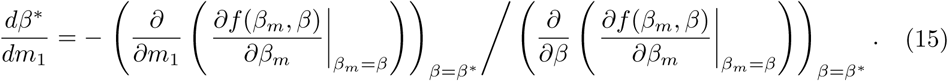

Because both ‘CSS’ and ‘EBP’ are convergence stable, which means that the denominator of equation (15) is always negative. Therefore, the sign of *dβ*^*∗*^*/dm*_1_ is determined by the numerator of equation (15). By direct calculations, the numerator of equation (15) is given by

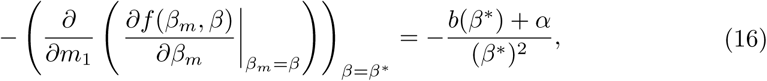

which is always negative when condition (2) holds. Therefore, for both the continuously stable strategy (CSS) and evolutionary branching point (EBP) *β*^*∗*^, *dβ*^*∗*^*/dm*_1_ *>* 0 is always true, that is to say, the evolutionarily singular strategy *β*^*∗*^ will always increase as the parameter *m*_1_ increases. This means that if the strength of density-dependent mortality *m*_1_ becomes strong, then the susceptible host population will increase its birth rate *b*(*β*) (because the transmission rate *β* will increase) and accordingly decrease its resistant ability, no matter the exact relationship between *b*(*β*) and *β*.

**Fig 5.**
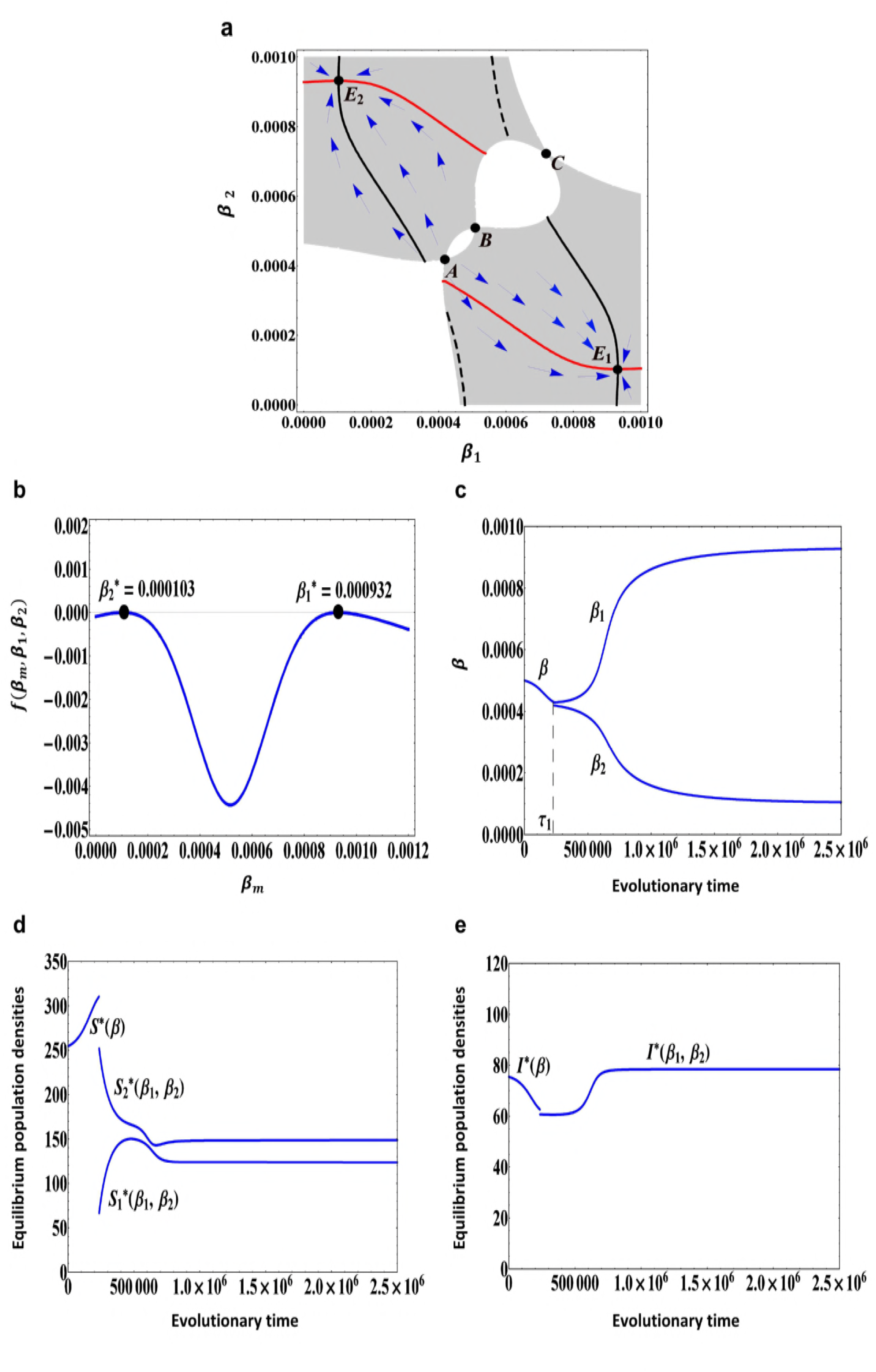
Bifurcation diagram of evolutionary dynamics when the demographic parameters are changed. (a) Evolutionarily singular strategy *β*^*∗*^ versus the pathogeninduced mortality rate *α* when the trade-off function is the same as in Fig.1a. (b) Evolutionarily singular strategy *β*^*∗*^ versus the natural death rate *m*_0_ when the trade-off function is the same as in Fig.1a. (c) Evolutionarily singular strategy *β*^*∗*^ versus the strength of density-dependent mortality *m*_1_ when the trade-off function is the same as in Fig.1a. (d) Evolutionarily singular strategy *β*^*∗*^ versus the pathogen-induced mortality rate *α* when the trade-off function is the same as in Fig.3a. (e) Evolutionarily singular strategy *β*^*∗*^ versus the natural death rate *m*_0_ when the trade-off function is the same as in Fig.3a. (f) Evolutionarily singular strategy *β*^*∗*^ versus the strength of density-dependent mortality *m*_1_ when the trade-off function is the same as in Fig.3a. In (a), (b), (c), (d), (e) and (f), the arrows indicate the directions of evolutionary change. The black solid curves indicate the evolutionarily singular strategies which are both convergence stable and evolutionarily stable; the red dashed curves indicate the evolutionarily singular strategies which are not evolutionarily stable. *CSS* denotes the continuously stable strategy, *EBP* denotes the evolutionary branching point. The grey region indicates combination of *β* and *b*(*β*) for which the positive population equilibrium is globally asymptotically stable. In (a), (b) and (c), other parameter values are the same as in Fig.1. In (d), (e) and (f), other parameter values are the same as in Fig.3.

Two typical examples are depicted in Figs.5c and 5f. Based on the globally concave trade-off function (12), we obtain the bifurcation diagram of evolutionarily singular strategy *β*^*∗*^ versus the mortality strength *m*_1_ (see Fig.5c). From Fig.5c, we can see that if the strength of density-dependent mortality *m*_1_ gradually increases, then the continuously stable strategy (CSS) *β*^*∗*^ will always increase and the susceptible host population will evolve to a state at which it has a lower resistant ability but with a higher birth rate and come to a halt. In this way the evolution of host resistance may benefit the whole susceptible host population. Moreover, based on the concave-convex-concave trade-off function (13), we obtain the bifurcation diagram of evolutionarily singular strategy *β*^*∗*^ versus the mortality strength *m*_1_ (see Fig.5f). From Fig.5f, we can see that for fixed other parameter values evolutionary branching of host resistance is possible when 0.00013 *< m*_1_ *<* 0.000144. Particularly, it can be seen that if *m*_1_ = 0, that is, there is no density-dependent mortality, then the evolutionary branching in the host resistance can not occur. If the strength of density-dependent mortality *m*_1_ gradually increases, then the value of evolutionary branching point (EBP) will increase and the susceptible hosts will firstly evolve to a state at which it has a lower resistant ability but with a higher birth rate and then branches into two different types. However, if the strength of density-dependent mortality *m*_1_ becomes relatively strong, then numerical simulation analysis shows that the evolutionarily singular strategy regains the evolutionary stability and becomes a ‘CSS’ (see Fig.5f).

Similarly, we examine the influence of the natural death rate *m*_0_ and the pathogeninduced mortality rate *α* on the continuously stable strategy (CSS) and evolutionary branching point (EBP) of host resistance. By the same argument as above, we can see that for both ‘CSS’ and ‘EBP’, *dβ*^*∗*^*/dm*_0_ *>* 0 and *dβ*^*∗*^*/dα >* 0, which means that the value of continuously stable strategy or evolutionary branching point *β*^*∗*^ will gradually increase as the parameter *m*_0_ or *α* increases (see Figs.5a, 5b, 5d and 5e). In particular, from Fig. 5d and Fig.5e, we can see that when parameters *m*_0_ and *α* take the middle range values, the evolutionary branching in the host resistance is possible. When the parameter *m*_0_ or *α* increases, the susceptible host population will firstly evolve to a state at which it has a lower resistance ability but with a higher birth rate and then branches into two different types. This means that in order to determine whether the evolutionary branching occurs, the susceptible host population must balance the costs and benefits based on the resistance ability, reproductive ability and other demographic conditions [10].

By the above analysis, we can see that if the constraint function is weakly convex in the vicinity of evolutionarily singular strategy, i.e., there is a weakly decelerating cost near the singular strategy *β*^*∗*^, then evolutionary branching in the host resistance is possible. From the bifurcation diagrams (see Figs.4 and 5), it can be seen that for a given trade-off function the evolutionary branching behavior can occur over a large range of demographic parameter values. After branching has occurred in the host resistance strategy, it becomes more interesting to study the further coevolution of such a dimorphic susceptible host. Next, we use a similar approach to investigate the evolutionary dynamics of such a dimorphic susceptible host and find the final evolutionary outcome.

## Results of dimorphic evolutionary dynamics

In this section, we extend the analysis to host population with two resident resistance strategies *β*_1_ and *β*_2_, and investigate whether the two types of susceptible hosts with different resistant abilities can evolve to an evolutionarily stable equilibrium at which they can stably coexist on the much longer evolutionary timescale [53, 54]. We assume that the dimorphic susceptible host shares all the properties except for resistant ability and natural birth rate. Moreover, it is assumed that mutations are very rare and very small.

### Coevolutionary model

Once the susceptible host population branches into two different types, *S*_1_ and *S*_2_, one with resistance strategy *β*_1_ and the other with a slightly different resistance strategy *β*_2_, then the population dynamics is given by

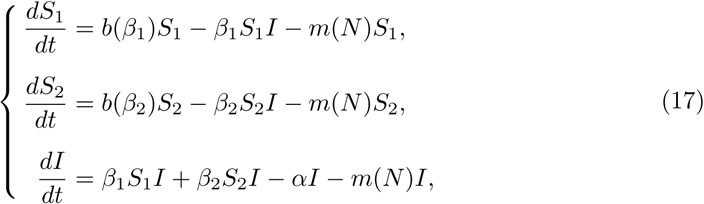

Where

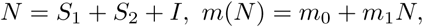

and *b*(*β*) is a monotonically increasing function with respect to *β* and eventually approaches to a state of saturation. For model (17), when the following condition (18) is true

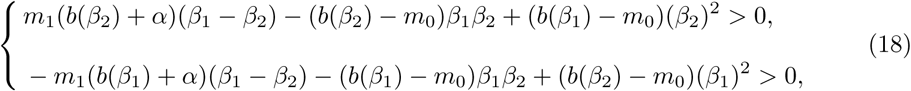

we obtain a globally asymptotically stable equilibrium 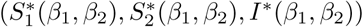 (see S3 Appendix for a detailed mathematical proof), which is given by

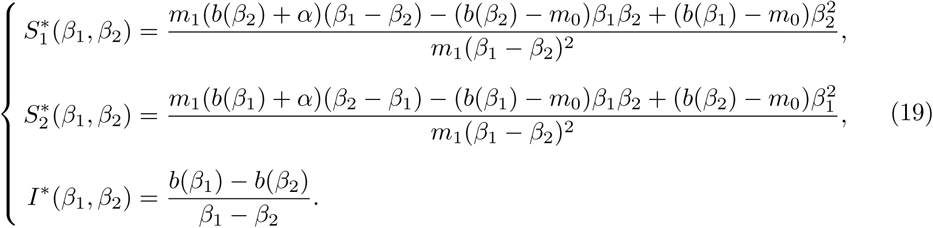

Because of the rarity of mutations, we assume that there is either a mutant host arising from susceptible host *S*_1_ or a mutant host arising from susceptible host *S*_2_, but not both at a time. By the same demonstrations as above, when a mutant susceptible host with a slightly different resistance strategy *β*_*m*_ enters into the resident community with a low density, the invasion fitness for the mutant host is then given by

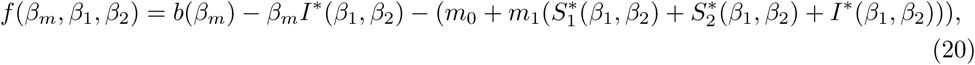

where *b*(*β*_*m*_) is the trade-off function and 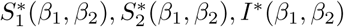 are respectively the equilibrium population densities of susceptible host *S*_1_, susceptible host *S*_2_ and infected host, which are described as in (19).

By using the same method as in S2 Appendix, we can rigorously demonstrate that if *f* (*β*_*m*_, *β*_1_, *β*_2_) *>* 0, then the mutant susceptible host will invade and cause a trait substitution. The evolutionary direction of host resistance is determined by the sign of selection gradients *g*_1_(*β*_1_, *β*_2_) and *g*_2_(*β*_1_, *β*_2_), which are given by

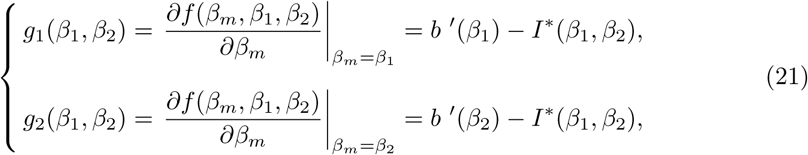

where *b* ^*′*^(*β*_*i*_) represents the first derivative of *b*(*β*_*m*_) with respect to *β*_*m*_ evaluated at *β*_*m*_ = *β*_*i*_, (*i* = 1, 2).

Furthermore, if the mutations are of small effect and rare, then the coevolutionary dynamics of host resistance is given by [30]

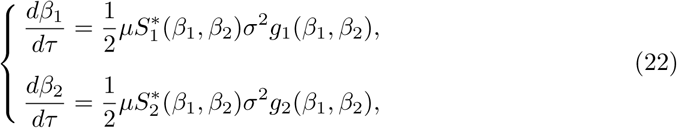

where time *τ* spans the evolutionary timescale and *μ* denotes the probability that a birth event in susceptible host *S*_*i*_ is a mutant, (*i* = 1, 2). 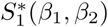 and 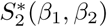 are the equilibrium population densities of the two susceptible hosts which are described in (19). *σ*^2^ denotes the variance of phenotypic effect of a mutation in susceptible host *S*_*i*_, (*i* = 1, 2). *g*_1_(*β*_1_, *β*_2_) and *g*_2_(*β*_1_, *β*_2_) are the selection gradients described in (21). Because we assume that the two host residents differ in only their birth rates and their resistance, they have the same *μ* and *σ*^2^. Moreover, we also assume that these two parameters do not depend on the trait values and are constants, therefore parameters *μ* and *σ*^2^ are the same as in the monomorphic host population. Model (22) tells us how the expected values of host resistance will change after evolutionary branching [53, 54].

### Evolutionarily singular dimorphism

We firstly identify the specific feature of a trade-off function that can support an evolutionarily singular dimorphism. The directional evolution may come to a halt at an evolutionarily singular dimorphism 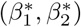. From (21), we can see that the condition for a positive singular dimorphism 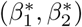 is given by

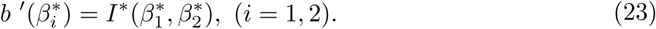

From (23), we can see that a trade-off function that can support a singular dimorphism must have two points at which it has the same slope, which cannot be the case for a globally concave trade-off function (such as (12)). If a singular dimorphism does not exist, then in simple trade-offs two host strains may continue to coexist indefinitely at the respective maximum and minimum trait values. By inserting (19) into (23), we obtain a specific expression for the required slope

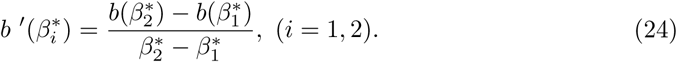

Therefore, a necessary condition for *b*(*β*) to support a positive singular dimorphism is that it is tangential to a straight line at two separate points [28, 64]. The simplest case is a concave-convex-concave trade-off function.

### Continuously stable coexistence

If the singular dimorphism 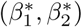 is both convergence stable and evolutionarily stable, it is called a continuously stable dimorphism and it represents the finally evolutionary outcome, at which the dimorphic susceptible host can stably coexist on the long-term evolutionary timescale [28, 53, 54, 64].

Specifically, the convergence stability of 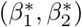 can be seen from the Jacobian matrix of system (22) at the singular dimorphism [61, 63]. Using (24), the Jacobian matrix *J*_1_ of system (22) at 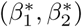 is given by

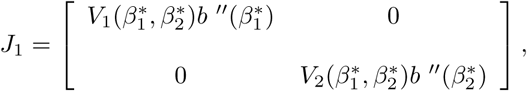

where 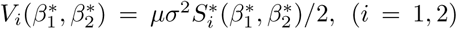. 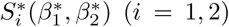 are described in (19) with 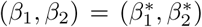. We can see that if 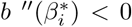 for *i* = 1 and 2, then the determinant of Jacobian matrix *J*_1_ is positive and its trace is negative. Therefore, if the trade-off function is concave at both 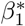 and 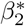, then the evolutionarily singular dimorphism 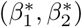 is locally convergence stable [28, 53, 54, 64].

Besides, if

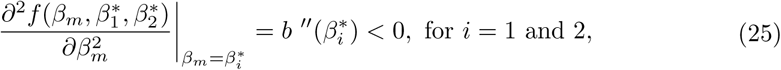

then the singular dimorphism 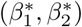 is locally evolutionarily stable. This condition is satisfied if the trade-off function is concave at both 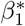 and 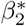. This means that once a positive singular dimorphism exists and is convergence stable, then it is always evolutionarily stable, and no further evolutionary branching is possible [28, 53, 54, 64]. Therefore, we obtain the following result.

**Theorem 3** *Assuming condition (18) holds*. *If condition (24) is satisfied and 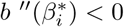 for i* = 1 *and* 2, *then the evolutionarily singular dimorphism 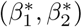 of system (22) is continuously stable*.

From Theorems 2 and 3, we can see that if there is a weakly decelerating cost near the singular strategy, then evolutionary branching in the host resistance will occur. After branching, for a concave-convex-concave trade-off function, there may exist a continuously stable singular dimorphism, the susceptible host population will finally evolve into two different types, which can stably coexist on the much longer evolutionary timescale [28, 64]. In this case, the final evolutionary outcome contains two types of hosts with different susceptibility.

**Fig 6.**
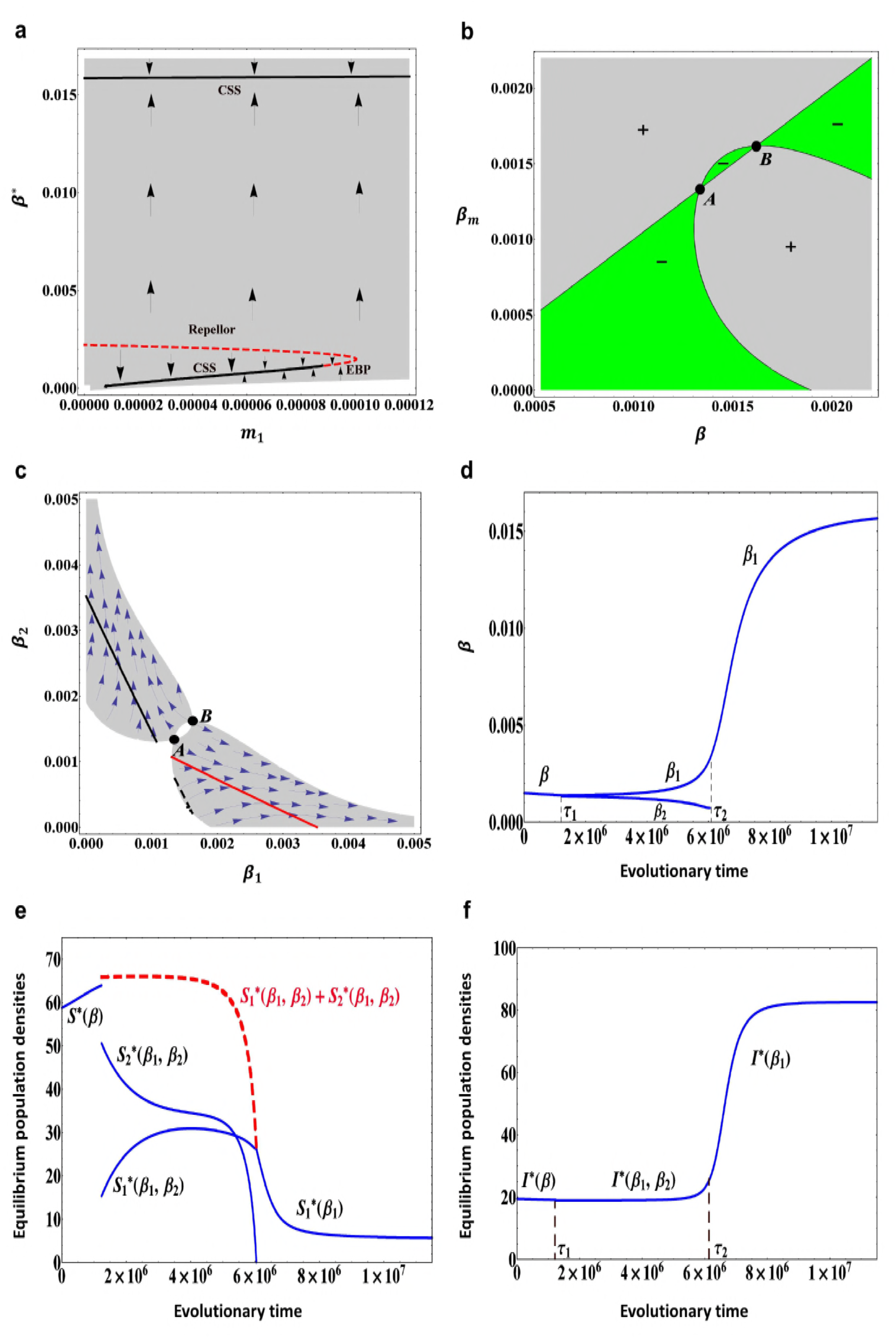
Dimorphic evolutionary dynamics of host resistance when the tradeoff function is given by (13) with parameters. *b*_0_ = 0.055, *b*_1_ = 4.0, *b*_2_ = 0.05, *b*_3_ = 0.05, *β*_0_ = 0.0005, *σ*_*c*_ = 0.0002. (a) A trait evolution plot. The singular strategy *A* = 0.000419 is an evolutionary branching point. After the branching has occurred, there exist two dimorphic singular strategies *E*_1_ = (0.000932, 0.000103) and *E*_2_ = (0.000103, 0.000932). The vector fields obtained from the deterministic model (22) indicate the directions of evolutionary change of traits *β*_1_ and *β*_2_. The black curves and red curves indicate the isoclines of traits *β*_1_ and *β*_2_, respectively. The solid curves indicate the evolutionarily singular strategies which are evolutionarily stable, whereas the dashed curves indicate the evolutionarily singular strategies which are not evolutionarily stable. (b) A fitness landscape plot when 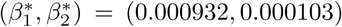. (c) A simulated evolutionary tree obtained through simulation of models (6) and (22) with the initial condition *β*(0) = 0.0005. At evolutionary time *τ*_1_ evolutionary branching occurs. (d) Equilibrium population densities of susceptible hosts when the traits *β*_1_ and *β*_2_ evolve. (e) Equilibrium population density of infected host when the traits *β*_1_ and *β*_2_ evolve. Other parameter values: *α* = 0.075, *m*_0_ = 0.006, *m*_1_ = 0.00014, *μ* = 0.01, *σ* = 0.00001.

As an example, we take the same trade-off function and parameter values as in Fig.3. In this case, evolutionary branching in the host resistance is possible (see Figs.3b and 3c). After branching has occurred in the host resistance, the two types of susceptible hosts evolve according to model (22). By numerical simulation, we find two evolutionarily singular dimorphisms *E*_1_ = (0.000932, 0.000103) and *E*_2_ = (0.000103, 0.000932) (depending on the initial trait value, only one of the two singular dimorphisms can be reached). At the two singular strategies we find that 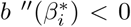 for *i* = 1 and 2, thus the two evolutionarily singular dimorphisms are continuously stable. Fig.6a shows that the evolutionarily singular dimorphisms are convergence stable and Fig.6b shows that the evolutionarily singular dimorphism *E*_1_ = (0.000932, 0.000103) is evolutionarily stable. Therefore, after branching in the host resistance has occurred, depending on the initial resistant ability, the dimorphic susceptible host will evolve towards one of the two singular dimorphisms *E*_1_ and *E*_2_ and come to a halt. The final evolutionary outcome contains two types of susceptible hosts, which can stably coexist on a long-term evolutionary timescale.

The evolutionary tree of host resistance obtained through simulation of models (6) and (22) with initial trait value *β*(0) = 0.0005 is depicted in Fig.6c. At evolutionary time *τ*_1_, the susceptible host branches into two different types. After branching has occurred, we can see that the trait in one branch initially increases, whereas in the other branch it initially decreases. This was expected because at a branching point the resident and mutant susceptible host coexist under opposite selection pressure and diverge in their resistant strategies [28, 42, 53, 54, 64]. Finally, the susceptible host evolves into two different types, one is a relatively higher susceptible strain, the other is a relatively higher resistant strain. The corresponding equilibrium population densities of susceptible hosts and infected host along the evolutionary tree are depicted in Figs.6d and 6e. It can be seen that the equilibrium population densities of the two types of susceptible hosts differ very much just after the branching. The susceptible host *S*_1_ associated with the upper branch has a lower equilibrium density, but its population equilibrium density initially gradually increases as its trait evolves. This maybe because with the increase of transmission rate *β*_1_ (i.e., with the decrease of resistant ability), the birth rate of susceptible host *S*_1_ will also increase, the advantage far outweighs its disadvantage. By comparison, the susceptible host *S*_2_ associated with the lower branch has a higher equilibrium density, but its equilibrium density initially gradually decreases as its trait evolves (the equilibrium densities immediately after branching depend on how the initial trait values of the dimorphic population are chosen; a different initial point in the vicinity of the branching point may lead to the opposite results). Finally, at the evolutionarily stable dimorphism *E*_1_ = (0.000932, 0.000103) the equilibrium population densities of the two types of susceptible hosts are very close to each other. From Fig. 6e, we can also see that after branching has occurred, the equilibrium population density of infected host will gradually increase and finally reach to a saturation state. In other words, the dimorphic evolution of susceptible host increases the equilibrium population density of infected host. On the whole, for a concave-convex-concave type of trade-off curve, under certain conditions the dimorphic susceptible host can continuously stably coexist on a long-term evolutionary timescale [53, 54].

### Evolutionary extinction of one emerging branch

From Theorem 2 and bifurcation analysis in Section 3.3 and Section 3.4, we can see that the evolutionary branching in the host resistance is possible for many different types of trade-off functions and under different demographic conditions. However, the evolutionary branching does not ensure that the dimorphic susceptible host can continuously stably coexist on a long-term evolutionary timescale. The further coevolution of the dimorphic susceptible host may result in loss of one or two evolutionary branches and thus the dimorphic susceptible host will go back to a monomorphism or become extinct [28, 42, 53, 54]. In the present model, we find that after branching has occurred in the host resistance, the condition (24) in Theorem 3 may not be met, thus there may not exist a positive singular dimorphism in system (22). In this case, the dimorphic susceptible host can not persistently coexist on the long-term evolutionary timescale. Through numerical simulation analysis, we find that after the branching has occurred, depending on the specific shape and strength of trade-off relationship between host resistance and its fertility, one of the two evolutionary branches may become extinct, and the dimorphic susceptible host finally falls back to a monomorphism [28, 42, 53, 54]. However, it’s worth noting that it is not necessary that the two host strains will evolve outside of the coexistence region whenever there is no dimorphic singular point. Depending on the evolutionary speed and trade-off function, they may just sit at their maxima and minima provided there is a route through the coexistence region.

As an example, we take the same trade-off function as in (13). However, the parameter values are assumed to be *b*_0_ = 0.05, *b*_1_ = 5.5, *b*_2_ = 0.05, *b*_3_ = 0.8, *β*_0_ = 0.0018 and *σ*_*c*_ = 0.008. In this case, the trade-off function *b*(*β*) in (13) is monotonically increasing and finally converges to a saturation state, which looks like a sigmoidal curve. Other parameter values are *α* = 0.075, *m*_0_ = 0.0055, *μ* = 0.01, *σ* = 0.00005. The bifurcation diagram of evolutionarily singular strategy *β*^*∗*^ versus the strength of density-dependence mortality *m*_1_ is depicted in Fig.7a. We can see that when 0.000089 *< m*_1_ *<* 0.000101, the smaller singular strategy is convergence stable but not evolutionarily stable, thus evolutionary branching in the host resistance is possible. The pairwise invasibility plot is shown in Fig.7b when *m*_1_ = 0.000098, which indicates the evolutionarily singular strategy *A* is an evolutionary branching point (EBP). After branching has occurred in the host resistance, the two types of susceptible hosts evolve according to model (22). However, in this case, by numerical simulation analysis we do not find any positive evolutionarily singular dimorphism when 0.000089 *< m*_1_ *<* 0.000101 (see Fig.7c). Therefore, after branching has occurred, the dimorphic susceptible host may evolve towards the edge of coexistence region (see Fig.7c). Near the edge of coexistence region, the next successful mutant host will drive the population outside the coexistence region, so that the dimorphic susceptible host will go back to a monomorphism. In particular, we find that independent of the initial resistant ability, always the evolutionary branch with the lower *β* goes extinct and the remaining branch with higher *β* will continue to evolve to a continuously stable strategy (CSS). In this case, the final evolutionary outcome contains a monomorphic susceptible host with a relatively weaker resistance, but it has a relatively higher birth rate. The evolutionary tree of host resistance obtained through simulation of models (6) and (22) with initial trait value *β*(0) = 0.0015 is depicted in Fig.7d. At evolutionary time *τ*_1_, the susceptible host population experiences evolutionary branching. After branching, we can see that the trait in one branch initially increases and finally evolves to a continuously stable strategy, while in the other branch it initially decreases and at evolutionary time *τ*_2_ it becomes extinct. This phenomenon can be seen as an typical example of ‘evolutionary murder’ [28,39–43,54]. Particularly, from Figs.7b, 7c and 7d, we can see that even if all mutations are small, the combination of evolutionary branching and loss of one emerging branch has efficiently allowed the initially monomorphic susceptible host to jump across the repellor *B* and to continue its monomorphic evolution on the other side [28, 39–43].

**Fig 7.**
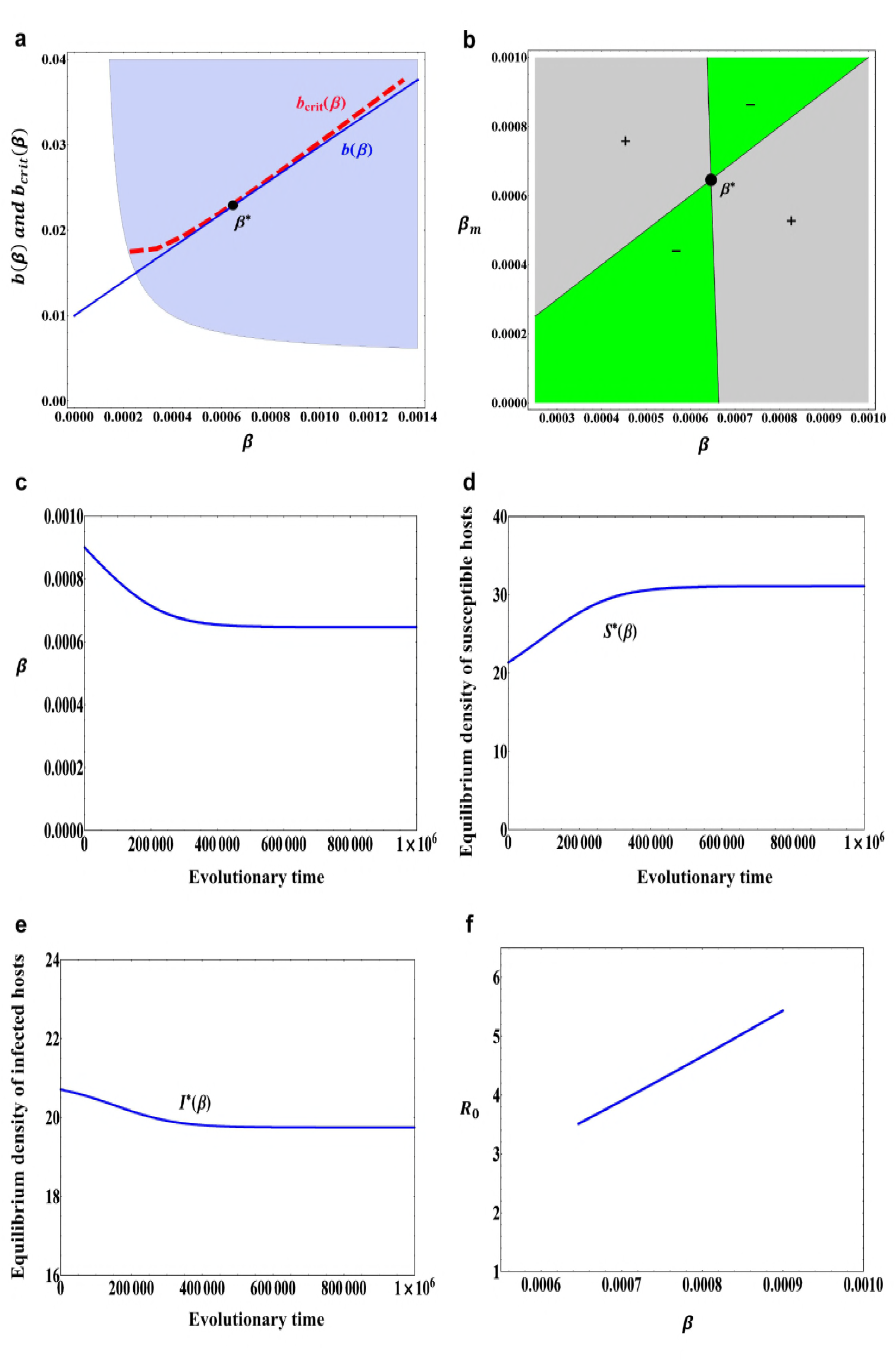
Evolutionary extinction of one emerging branch when the trade-off function is given by (13) with parameters. *b*_0_ = 0.05, *b*_1_ = 5.5, *b*_2_ = 0.05, *b*_3_ = 0.8, *β*_0_ = 0.0018 **and** *σ*_*c*_ = 0.008. (a) Evolutionarily singular strategy *β*^*∗*^ versus the strength of density-dependent mortality *m*_1_. Arrows indicate the directions of evolutionary change. *CSS* denotes the continuously stable strategy, *EBP* denotes the evolutionary branching point. (b) A pairwise invasibility plot when *m*_1_ = 0.000098. (c) A trait evolution plot when *m*_1_ = 0.000098. The vector fields obtained from the deterministic model (22) indicate the directions of evolutionary change of traits *β*_1_ and *β*_2_. The black curves and red curves indicate the isoclines of traits *β*_1_ and *β*_2_, respectively. The solid curves indicate the evolutionarily singular strategies which are evolutionarily stable, whereas the dashed curves indicate the evolutionarily singular strategies which are not evolutionarily stable. (d) A simulated evolutionary tree obtained through simulation of models (6) and (22) with initial condition *β*(0) = 0.0015 and *m*_1_ = 0.000098. At evolutionary time *τ*_1_ evolutionary branching occurs and at evolutionary time *τ*_2_ the emerging branch *β*_2_ become extinct. (e) Equilibrium population densities of susceptible hosts when the traits *β*_1_ and *β*_2_ evolve. At evolutionary time *τ*_2_, the equilibrium population density of susceptible host *S*_2_ becomes zero. The dashed curve indicates the total equilibrium population density of the two types of susceptible hosts. (f) Equilibrium population density of infected host when the traits *β*_1_ and *β*_2_ evolve. Other parameter values: *α* = 0.075, *m*_0_ = 0.0055, *μ* = 0.01, *σ* = 0.00005.

The corresponding equilibrium population densities of susceptible hosts and infected host along the evolutionary tree are depicted in Figs.7e and 7f. We can see that the equilibrium population densities of the two types of susceptible hosts differ very much just after the branching and thereafter the equilibrium population density of susceptible host *S*_2_ gradually decreases. At time *τ*_2_, due to the trade-off relationship between host resistance and its fertility, the equilibrium population density of susceptible host *S*_2_ becomes zero [28, 42, 53, 54]. Finally, the susceptible host *S*_1_ evolves to a continuously stable strategy, at which it has a relatively weaker resistance but with a relatively higher birth rate. Especially, from Figs.7e and 7f, we can see that the evolution of host resistance gradually reduces the equilibrium population density of susceptible hosts, on the contrary it gradually increases the equilibrium population density of infected host.

## Discussion and conclusions

Understanding the evolutionary dynamics of host resistance in the light of ecological feed-backs has important implications for the prevention and control of infectious disease. In this paper, based on a classical susceptible-infected (SI) model and the method of adaptive dynamics, we investigated the evolutionary mechanism of host resistance to pathogen infection. We assumed that only the resistance-related trait of host population can adaptively evolve, but there exists a trade-off relationship between the host resistance and its fertility. By using the technique of critical function analysis, we identified the general features of trade-off function that allow for a continuously stable strategy and evolutionary branching of host resistance. We found that the final evolutionary outcomes depend mainly on the shape and strength of trade-off function. In other words, which evolutionary outcome will occur relies largely on a balance between costs (here, reduction in birth rate) and benefits (here, resistance to transmission of the pathogen) [10, 13, 28]. We showed that if there is an accelerating cost (i.e., the trade-off curve is concave) when the host resistance evolves, then a continuously stable strategy is predicted. In particular, for a globally concave trade-off function, we found that if the initial resistant ability is relatively strong (*β*(0) = 0.0003), then the evolution may reduce the resistant ability of susceptible host (i.e., in order to increase the birth rate of susceptible host), the equilibrium population density of susceptible host may gradually decrease, whereas the equilibrium population density of infected host may gradually increase and finally reach to a saturation state. In this case, the evolution of host resistance may lead to an increase of basic reproduction number *R*_0_. However, if the initial resistant ability is relatively weak (*β*(0) = 0.0009), then the evolution of host resistance may result in a decrease of basic reproduction number *R*_0_. In general, *R*_0_ is almost never maximized [65]. Alternatively, with a weakly decelerating cost (i.e., the trade-off curve is weakly convex) and density-dependent mortality, we found that evolutionary branching in the host resistance is possible [10, 13, 28, 53, 54].

Moreover, by using the method of implicit differentiation, we found that independent of the trade-off function, the values of continuously stable strategy (CSS) and evolutionary branching point (EBP) will always increase as the strength of density-dependent mortality *m*_1_ or the natural death rate *m*_0_ or the pathogen-induced mortality rate *α* increases. Through evolutionary bifurcation analysis, it was also found that evolutionary branching in the host resistance is possible for a wide range of values of demographic parameters and different types of trade-off functions. Particularly, the presence of a density-dependent mortality *m*_1_ plays a crucial role in the evolutionary branching of host resistance. If there is no density-dependent mortality, then the two different types of susceptible hosts can not coexist in the population community (because in this case there is no positive population equilibrium for model (17)), thus evolutionary branching in the host resistance is impossible [10, 13, 28, 45, 47, 53, 54]. This means that in order to determine whether the branching occurs, the susceptible host must balance the costs and benefits on the basis of the resistance ability, reproductive ability and other demographic conditions. It should be noted that in this paper we only considered a simple susceptible-infected (SI) model. It would be interesting to study the phenomenon of evolutionary branching in the host resistance based on a more realistic epidemic model, such as susceptible-infected-susceptible (SIS) model or susceptible-infected-recovered (SIR) model. In addition, defence mechanisms in hosts include not only avoiding becoming infected, but also recovering more quickly after infection or surviving longer once infected, therefore, it is also interesting to explore the evolution of costly host resistance with different defence mechanisms, such as recovery (increase in the clearance rate) or tolerance (reduction in the death rate due to infection) [10, 13, 28, 45, 47].

After evolutionary branching in the host resistance has occurred, we found that depending on the evolutionary speed and the specific shape and strength of trade-off relationship, the susceptible host population may evolve into an evolutionarily stable dimorphism or the evolutionary branching is followed by the loss of one host strain. For one thing, if the trade-off curve is concave-convex-concave and tangential to a straight line at two separate points, then for a large range of demographic parameter values the susceptible host population will eventually evolve into two different types, which can continuously stably coexist on the much longer evolutionary timescale [28, 53, 54, 64]. Numerical simulation analysis furthermore showed that the equilibrium population densities of the two types of susceptible hosts are very close to each other finally (whether this is a general phenomenon needs to further study), and the dimorphic evolution of susceptible host may increase the equilibrium population density of infected host. Therefore, we can say that for a type of concave-convex-concave trade-off relationship, in the simple susceptible-infected community the monomorphic susceptible host can become dimorphic and the dimorphic susceptible host can continuously stably coexist on the long-term evolutionary timescale, no further evolutionary branching is possible [28, 53, 54, 64]. However, it should be noted that such coexistence is not necessarily stable against other life-history traits evolving on a much longer evolutionary timescale (such as recovery rate and pathogenicity) [28, 44, 54]. Therefore, it remains interesting to investigate how robust the coexistence is and under what conditions the evolutionary branching in the host resistance can induce the secondary branching in the infected host population.

For another, if the trade-off function looks like a sigmoidal curve and there is no straight line which is tangential to the trade-off curve at two separate points, then the evolutionary branching of host resistance may be possible, but it can not bring about an evolutionarily stable dimorphism. On the contrary, after branching has occurred, depending on the initial resistant ability, the dimorphic host will firstly evolve to the edge of coexistence region, and then the evolutionary branch with lower transmission rate will become extinct (i.e., the corresponding equilibrium population density will become zero) and the remaining branch will continue to evolve to a continuously stable strategy and come to a halt. This phenomenon is called as ‘evolutionary murder’ [28, 39–43, 54], which implies that the dimorphism of susceptible host may be non viability on the long-term evolutionary timescale. From these two examples, we can see that for the same trade-off function, different shape and strength may lead to entirely different evolutionary outcomes. Our results further highlights the importance of trade-offs to evolutionary behaviour of host resistance [13, 28]. In particular, numerical simulations showed that the evolution of host resistance may reduce the equilibrium population density of susceptible hosts, in contrast it may increase the equilibrium population density of infected host. This phenomenon is a bit of a violation of our intuition and has not been observed in previous works. In addition, we also found that even though all mutations are small, the combination of evolutionary branching and loss of one host strain has effectively allowed the initially monomorphic host population to jump across the repellor and to continue its monomorphic evolution on the other side [28, 39–43, 47, 53, 54]. It’s worth noting that if a singular dimorphism does not exist, then depending on the evolutionary speed and trade-off function, two host strains may just sit at their maxima and minima provided there is a route through the coexistence region or they may continue to coexist indefinitely at the respective maximum and minimum trait values. Moreover, except for an interior singularity or extinction of one branch, a dimorphic host population is likely to evolve to a boundary singularity where one branch has trait valve *β* = 0. These are still interesting questions worth further studying.

In general, the novelty of this study is reflected in the following several aspects. Firstly, our model assume that both susceptible host and infected host are subject to density-dependent mortality, which is more reasonable in the real world. Based on this population dynamic model, we can further study whether the evolutionary branching in the host may lead to evolutionary branching in the pathogen. Secondly, the trade-off function we used is a monotonically increasing function and eventually converges to a saturation state, which is more aligned with empirical evidence [9, 10, 13, 17, 18, 22]. Particularly, we found that for the same trade-off function, the different shape and strength may lead to entirely different evolutionary outcomes of host resistance. Thirdly, we performed the bifurcation analysis of evolutionary dynamics and discussed the influence of demographic parameters on the evolutionary behaviour of host resistance, which is helpful for us to develop an analytical method for the bifurcation theory of evolutionary dynamics. Fourthly, for every step of evolutionary invasion analysis, by using the method of Lyapunov function we have all performed rigorously mathematical proofs, such as the stability of population equilibrium and invasion implies fixation. After evolutionary branching in the host resistance has occurred, we also derived the demographic and evolutionary conditions that allow for a continuously stable coexistence of the dimorphic host and found an example of the loss of one host strain. In another word, based on a simple susceptible-infected (SI) model, we found very rich evolutionary behaviour of host resistance under different trade-off relationships. Therefore, this paper also provided a typical example for the evolutionary invasion analysis of host resistance by using the method of adaptive dynamics.

## Supporting Information

**S1 Appendix. Global asymptotical stability of** (*S*^*∗*^(*β*), *I*^*∗*^(*β*)).

**S2 Appendix. Invasion implies fixation.**

**S3 Appendix. Global asymptotical stability of 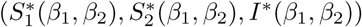.**

## Acknowledgments

We would like to thank Profs. Yicang Zhou and Yanni Xiao for their valuable discussion. The study was supported by grants from the National Natural Science Foundation of China (11571272 and 11631012), grant from the National Science and Technology Major Project of China (2012ZX10002001), grant from the Natural Science Foundation of Shaanxi Province (2015JQ1011), and grant from the China Postdoctoral Science Foundation (2014M560755).

